# Dueling Endogenous Viral-Like Sequences Control Synaptic Plasticity

**DOI:** 10.1101/2023.12.20.572708

**Authors:** P. Githure M’Angale, Adrienne Lemieux, Yumeng Liu, Jasmine Graslie, Shuhao Wang, Alfred Simkin, Vivian Budnik, Brian A. Kelch, Travis Thomson

## Abstract

The function of a large part of most genomes, generally called “junk DNA”, remains largely unknown. Much of this enigmatic DNA corresponds to transposons, which are considered genomic parasites. Here, we show the protein of the Ty1 retrotransposon *Copia* is enriched at the *Drosophila* neuromuscular junction and is transported across synapses. Unexpectedly, disrupting *Copia* expression results in increases in both synapse development and structural synaptic plasticity. Plasticity is kept in balance as *Copia* antagonizes the *Drosophila Arc* (activity-regulated cytoskeleton-associated protein) homolog, which is a transposon-derived gene. Our cryo-EM structure of the *Copia* capsid shows a shell with large cargo capacity and leads to a hypothesis for mutual antagonism of Arc and *Copia* capsid assembly. Our findings provide evidence that a fully functional transposon plays a role at synapses, suggesting that transposons and other types of ‘junk DNA’ are essential to developmental and cellular processes.

## INTRODUCTION

Large portions of eukaryotic genomes have undescribed functions and have been historically referred to as “junk DNA.” A substantial portion of junk DNA is comprised of transposable elements (TEs), which are mobile genetic elements capable of moving positions within a genome. This ability to insert sequences into new locations is regarded as ‘parasitic’ or ‘selfish.’ Host organisms bear the consequences of TE insertion and activation, as both processes are coincident with and could be partially causal of some diseases (*1*). However, genomic TE insertions are also widely regarded as major drivers of evolution, and many “domesticated” TE-like sequences contribute to gene expression and/or function (*2, 3*). For example, the domestication of a TE-fragment gave rise to the gene Arc, whose product undergoes viral-like transfer across the neuromuscular junction (NMJ) to regulate synaptic plasticity in flies and has been shown to transfer between mammalian neuronal cells in culture (*4, 5*). There is evidence that suggests ‘junk DNA’ may be under a type of selective pressure (*6, 7*). Selective pressure on TEs may lead to domestication and the repurposing of proteins and regulatory sequences for novel molecular functions.

The viral-like packaging of *Drosophila* Arc (dArc1) in the synapse prompted us to look for other molecules that are transported across presynaptic compartments in this viral-like manner, and we discovered that *Copia* RNA is enriched in extracellular vesicles (EVs) (*5*). In the *Drosophila* genome, *Copia* is an abundant TE which contains a GAG region encoding capsid-like proteins analogous to those encoded in the genome of retroviruses. *Copia* has been extensively studied as a critical source of insight into the viral-like properties of TEs, as it was shown to have reverse transcriptase activity and form viral-like intermediates during replication (3–5). Here, we found the capsid protein of *Copia* is enriched at and transfers across *Drosophila* NMJs. Disrupting *Copia* expression at the NMJ leads to a striking increase in structural synaptic plasticity. Further, we observed that the Copia protein can auto-assemble into capsids. These capsids have a distinct structure from dArc1. We demonstrate that *Copia* and *dArc1* are in an antagonistic relationship phenotypically and genetically, which appears to be a mechanism to regulate the development of plasticity. Collectively, our observations reveal a novel role for *Copia* as a regulator of NMJ plasticity, placing *Copia* in a well-established plasticity pathway alongside *dArc1.* This novel form of TE domestication provides some of the first *in vivo* evidence that TEs play a direct and pivotal role in neuronal development.

## RESULTS

### *Copia^gag^* is enriched at and transfers across the Drosophila NMJ

Previously, we reported that a spliced form of *Copia* mRNA, *Copia^gag^*, and the capsid-like gene *dArc1* is enriched in EVs from *Drosophila* Schneider 2 (S2R+) cells (*5, 8*). *Copia^gag^* encodes the complete capsid and a small part of the POL region that contains the self-processing protease domain (Fig. 1A). Prior work on dArc1 mRNA and protein shows enrichment at the NMJ (*5*). Here, we generated a polyclonal antibody against Copia (α-Copia^Full^), which recognizes peptides encoded by the GAG and POL regions and can detect the full-length unspliced Copia peptide, and one against a peptide that is specifically encoded by the *Copia^gag^* spliced transcript (Fig. 1A, S1A). Western blots of tissues using α-Copia^Full^ revealed a 50 kDa isoform – which corresponds to the predicted size of Copia^gag^-in addition to the full-size protein (Fig. 1B). Probing with α-Copia^gag^ revealed that the spliced isoform is the predominant Copia species expressed in the larval central nervous system (CNS) (Fig. 1B, S1D). We carried out peptide competition assays to validate the specificity of the α-Copia^gag^ antibody to ensure only the protein product of the spliced transcript is recognized (Fig. S1B). The presence of peptides of varying sizes that bind to the α-Copia^Full^ antibody in the body wall muscle (BWM) tissue could be a product of viral peptide auto-cleaving, where full-length peptides are cleaved into functional small peptides (*9*), and is consistent with the viral nature of TEs such as *Copia* (*10, 11*).

**Figure 1.**
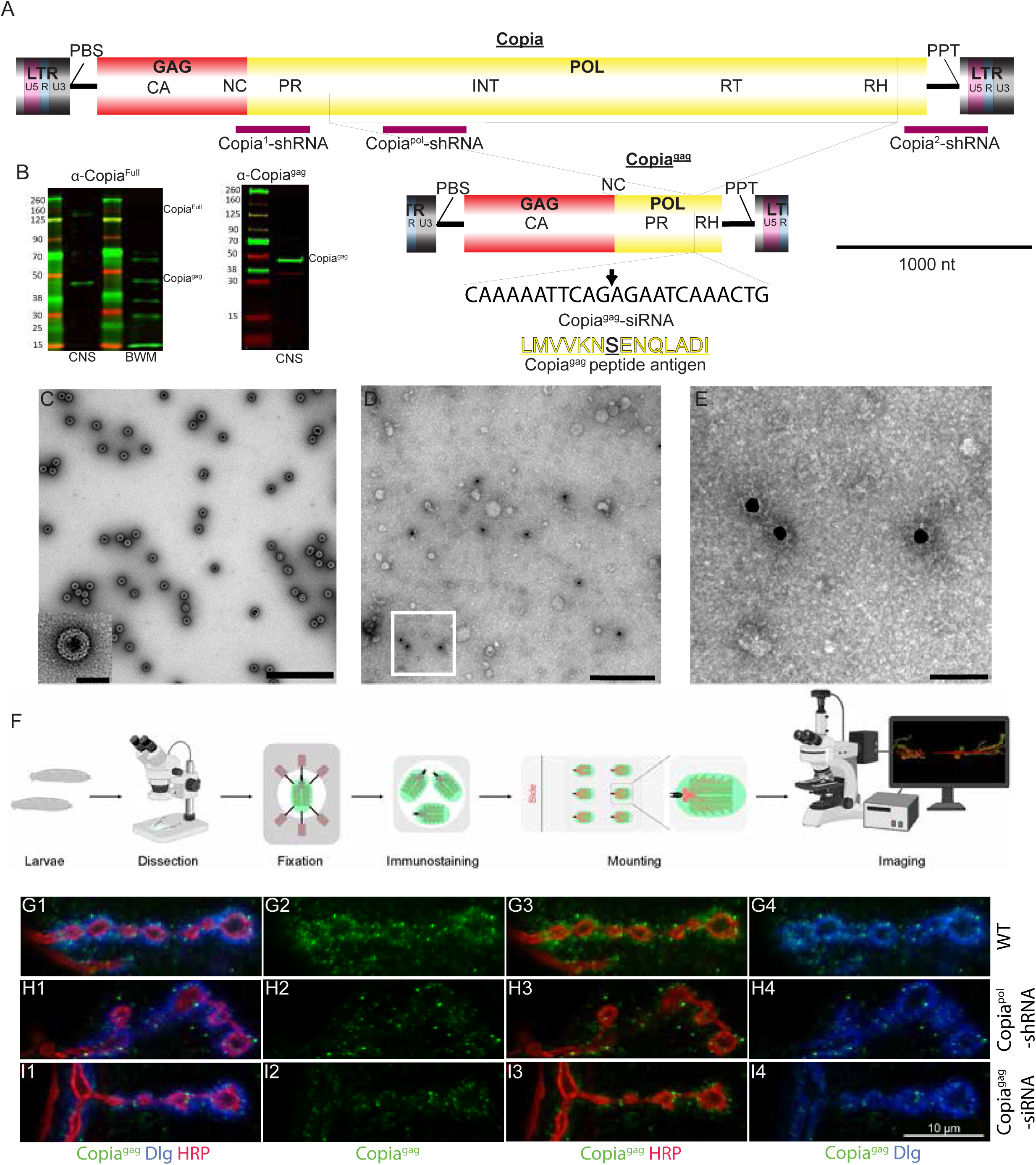
A spliced isoform of the retrotransposon, *Copia*, is enriched in neuronal tissue. A. Representation of the entire *Copia* genome (*Copia*^full^) and the spliced form (*Copia*^gag^). The location of the RNAi targets, splice site, and the peptide used to generate the Copia^gag^ antibody. B. Western blot of CNS lysate with the predicted size of Copia^gag^ indicated on the right and a molecular weight ladder on the left. C. Bacterially expressed Copia^gag^ self-assembles into capsid-like structures which are observable using negative stain EM, scale bar = 200nm; inset, close-up of an individual capsid, scale bar = 50nm. D. Representative EVs isolated from S2 cells were immuno-stained with α-Copia^gag^ (18nm gold secondary). E. A higher magnification of the closed square in D showing electron density of the α-Copia^gag^ stained S2 EVs (Scale bar = 50nm) F. Schematic of larval body wall muscle preparations for NMJ visualization. G. Representative confocal images of α-Copia^gag^ immunostaining show a striking enrichment at the wildtype *Drosophila* larval NMJ. H and I. Knockdown with either *Copia*^full^-shRNA (H) or *Copia*^gag^-siRNA (I) constructs expressed in the presynaptic compartment (C380-Gal4) cause a clear reduction of α-Copia^gag^ in both pre– and postsynaptic sides of the NMJ compared to wildtype (C380-Gal4/Canton-S). Abbreviations **-** Gag: Capsid forming proteins, which contains capsid (CA) and nucleocapsid (NC) sequences. Pol consists of the following subdomains: Protease (PR), Integrase (INT), reverse transcriptase (RT), and RNase H (RH). ENV: Envelope. The Long terminal repeat region (LTR) is needed for transposable element replication, consisting of U5, R, and U3 sub-regions. PBS: Primer Binding Site. PPT: Polypurine tract.

*Copia* has long been thought to form capsid-like structures in S2 cells, and we immunolabeled dense capsid-like structures with α-Copia^gag^ in detergent-treated EVs (Fig. 1D and 1E) (*9*). Capsid proteins form a shell-like structure that surrounds and protects the viral genome, and Copia^gag^ can form viral-like particles (VLPs) *ex vivo* (*12*) (Fig. 1C). We hypothesized that Copia^gag^ is acting in a viral-like manner and tested whether Copia^gag^ associates with the *Copia* transcript. We conducted RNA immunoprecipitations (RIP) from S2 cells as well as larval CNS and body wall muscles (BWM) tissues using α-Copia^gag^ and α-Copia^Full^ antibodies and probes for *Copia^gag^* and *Copia^full^* transcripts. We found that both α-Copia^Full^ and α-Copia^gag^ immunoprecipitated with the *Copia* transcript in CNS and BWM (Fig S1E), supporting a model that Copia forms capsids *in vivo* and encapsulates its own transcript, possibly along with transcripts of other genes.

Immunolabelling of *Drosophila* larvae with α-Copia^gag^ reveals a striking enrichment at the NMJ (Fig. 1F and 1G). To further confirm the specificity of the Copia antibodies, we leveraged the powerful Gal4/UAS system, which has been extensively used for precise repression or expression of genes in pre– and/or postsynaptic cells at the NMJ (*13*). We designed three UAS-shRNA constructs each targeting different regions of *Copia* to disrupt its expression (Fig. 1A). We observed a substantial reduction of α-Copia^gag^ signal when expressing *Copia^pol^*-shRNA using the motor neuron driver C380-Gal4 (Fig. 1G–I and S1H-K). We obtained a very similar effect with another shRNA construct, copia^1^-shRNA, which targets the GAG region that is distinct from copia^pol^-shRNA (Fig. S1H-I). We observed that expressing *Copia^pol^*-shRNA in neurons (see Fig. S1C for description and anatomy of boutons) caused a reduction of α-Copia^gag^ in the presynaptic boutons as well as in the postsynaptic muscle tissue (Fig. 1H). This strongly suggests that the postsynaptic localization of Copia^gag^ is at least partially derived from a pool of Copia mRNA and/or protein in the presynaptic cell (Fig. 1G-I). Due to the enrichment of α-Copia^gag^ signal at the larval NMJ, we designed a single siRNA construct, UAS-*Copia^gag^*-siRNA, that recognizes the *Copia^gag^* splice site to disrupt expression of only the spliced isoform, (Fig. 1A). Expressing UAS-*Copia^gag^*-siRNA resulted in a reduction of α-Copia^gag^ signal at the NMJ (Fig. 1G and 1I). We validated that disrupting *Copia* expression in larvae caused a significant decrease in *Copia* mRNA expression (Fig. S5A-B).

#### The *Copia* capsid structure reveals molecular basis for RNA encapsulation

To illuminate how Copia behaves in a virus-like manner, we sought to determine the structure of the *Copia* capsid. We purified capsids from the Copia^gag^ splice variant that we heterologously expressed in *E. coli*. The Copia^gag^ protein self-processes within *E. coli* to form separate Copia-GAG and Protease proteins (Fig. S4A and B). We isolated capsids of sufficient homogeneity that we could determine their structure using single-particle cryo-EM methods to ∼3.3-Å overall resolution (Fig. 2A, and Fig. S2 and S3). The *Copia* capsid adopts T=9 geometry with an inner radius of 220 Å. The Copia capsid is much larger than the T=4 dArc1 capsid (radius ∼170 Å), with ∼2.2-times increased internal capacity (Fig. 2B). There is a layer of featureless density lining the inner surface of the Copia capsid; we ascribe this density to the RNA that comigrates with the *Copia*-gag protein, as measured by A260/A280 ratio of ∼1.6. Because the RNA does not assume T=9 symmetry, its density is amorphous in our reconstruction. We hypothesize that the *Copia* capsid is larger than dArc1 because it packages larger cargo, such as the longer *Copia* mRNA (5 kbp vs 2 kbp), and the enzymes Reverse Transcriptase and/or Integrase.

**Figure 2.**
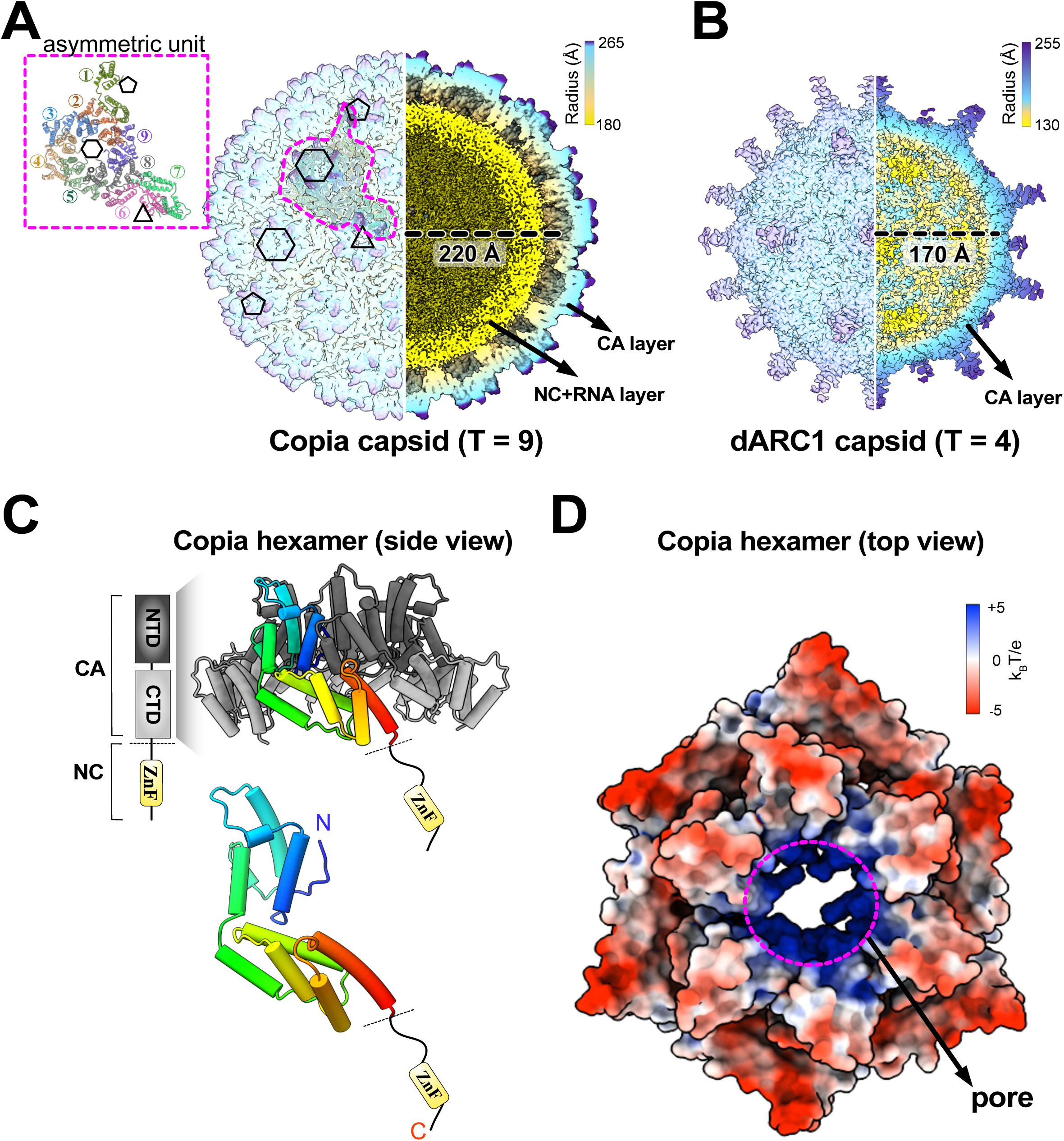
The cryo-EM structure of icosahedral Copia^Gag^ capsid. A. The electron density map of Copia^Gag^ capsid, colored by radius. Left half of the map is the surface of the capsid; right half of the map is a cross section view. The five-fold pentamer, three-fold hexamer and non-symmetric hexamers are denoted as black pentagon, triangle, and hexagons. The asymmetric unit is outlined in magenta dashed line on the map and shown in detail on the top left panel, highlighting the 9 monomers composing one asymmetric unit (T=9). The radius (220 Å) is measured between the center of the capsid and the mass center of the Copia^CA_CTD^ of the asymmetric unit. The outer layer is labeled as capsid (CA) layer, and the yellow inter layer density is labeled as nucleocapsid (NC) and RNA layer, the structure of which is undetermined due to disorder. B. The electron density map of dArc1 capsid colored by radius and determined in a previous study (*14*). The left half of the map is the surface of the capsid; right half of the map is a cross section view. The radius (170 Å) is measured between the center of the capsid and the mass center of the dArc1^CA_CTD^. The outer layer is labeled as capsid (CA) layer; the spikes on the surface of dArc1 capsid are unmodeled. There is no bound RNA in the assembled dArc1 capsid. C. The atomic model of Copia hexamer. Top panel, the model of CA region is colored with NTD in dark grey and CTD in light grey. The unstructured NC domain is shown as black line with the zinc finger domain (ZnF) located near the C-terminus. One monomer within the hexamer is colored in rainbow from N– to C-terminus. Bottom panel, the atomic model of one Copia^Gag^ monomer. D. Copia^Gag^ non-symmetric hexamer is colored by electrostatic potential. The positively charged (blue) pore is at the center of the hexamer.

The capsid layer consists of the Copia^gag^ protein’s CA region, which consists of two alpha-helical domains (Fig. 2C). The protein also contains the nucleocapsid or NC region, which harbors a Zn-finger RNA-binding domain. The NC region is not clearly visualized in the map but is inside the capsid and associated with the density attributed to RNA. The N-terminal and C-terminal domains of the CA region are structurally similar to the those seen in retroviruses and other retrovirus-like capsids (such as dArc1; C_α_ RMSD ∼ 3.5 Å and 2.7 Å for NTD and CTD, respectively) (*14*). However, the capsid structure is more similar to that of retroviruses than dArc1, because the *Copia* capsid lacks the spike emanating from the outer surface of the dArc1 capsid (*14*). In place of the spikes, the *Copia* capsid contains positively charged pores at the center of each hexameric or pentameric sub-assembly, similar to found in retroviruses such as HIV (*15*) (Fig. 2D). The positively charged pores of HIV act as gates for dNTP entry into the capsid for fueling reverse transcriptase activity (*16*). We hypothesize that the pores play a similar role in the *Copia* capsid.

We find that the *Copia* capsid formation requires RNA. If we remove the bound RNA using anion-exchange chromatography or by deleting the NC region, then we do not observe capsid assembly (Fig. S4C-E). However, the *Copia*^gag^-ΔNC variant can co-assemble with WT-*Copia* CA that carries RNA, which establishes the requirement for RNA (Fig. S4F). These results are distinct from dArc1, which can assemble in the absence of RNA (*14*), but similar to HIV, which requires RNA binding for efficient assembly (*17*). These results further support the proposal that *Copia* is functioning in a virus-like fashion.

#### *Copia* is a negative regulator of acute structural synaptic plasticity

Because Copia^gag^ forms virus-like particles at the NMJ similar to dArc1, we investigated whether it also plays a role in synaptic development and plasticity. Synaptic boutons at the NMJ are continuously formed throughout larval development, which correlates with larval muscle growth (*18*). Thus, a disruption of NMJ scaling is a measure of developmental synaptic plasticity (*19*). Expressing *Copia^pol^*-shRNA with the motor neuron-specific C380-Gal4 driver caused a striking ∼50% increase in synaptic bouton number (Fig. 3A, 3B, 3D, and S5E-F). This effect was also seen with the single siRNA construct directed against the *Copia^gag^* mRNA splice site (Fig. 3A, 3C, and 3D). In addition to an enhancement in the number of synaptic boutons, we also observed an increase in “hyperbudding,” which we define as the presence of three or more boutons budding off from a central, larger (parent) bouton (Fig. 3A3, 3B3, 3C3 and 3E).

**Figure 3.**
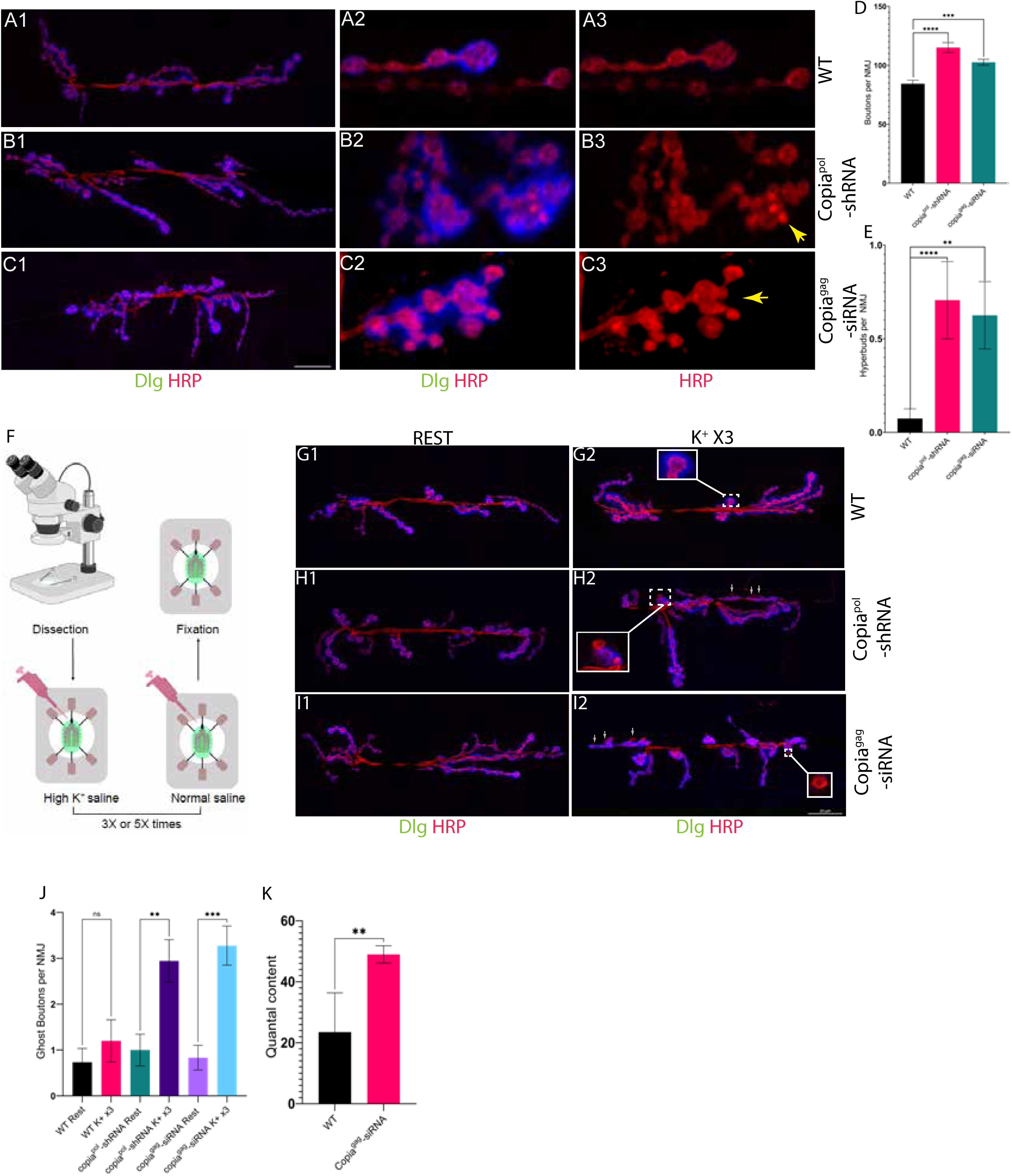
Disruption of *Copia* in motor neurons causes changes in synaptic development and plasticity. A –C. Expressing *Copia*^pol^-shRNA (B) or *Copia*^gag^-siRNA (C) constructs in the motoneurons of larvae causes increased bouton formation at NMJs compared to wildtype (C380-Gal4/Canton-S) (A). There is a substantial increase in hyperbudding (A3 vs. B3 and C3), where two or more boutons “bud” off the same central bouton (yellow arrows designate budding boutons in B3 and C3). D. Quantification of bouton numbers. E. Quantification of hyperbudding events. F. Schematic of activity paradigm for larvae stimulated with potassium. G–I. Stimulating wildtype flies (G) with three rounds of potassium treatments does not induce them to form new synapses (ghost boutons) but does induce increased bouton formation after disrupting expression of Copia (H and I) J. Quantification of data represented in G–I. K. NMJs with a presynaptic disruption of *Copia*^gag^ show an increase in synaptic vesicle release (quantal content). DLG = α-Discs Large (postsynaptic marker), HRP = α-horseradish peroxidase (presynaptic marker). N for D and E= from left to right; number of animals/NMJs quantified, 14/27, 9/17, 9/16, for J N= 12/19, 10/13, 6/12, 9/17, 6/11, 9/18. WT is C380-Gal4/Canton-S for panels A and G. ns p≥0.05, * p<0.05, ** p<0.01, *** p<0.001, and **** p<0.0001.

Our discovery that *Copia* is a negative regulator of bouton formation prompted us to test whether it also affects rapid activity-dependent bouton formation (*20*). Dissected larval NMJs can be acutely stimulated (High K^+^; 90 mM) to induce new synaptic bouton formation (*20*). These nascent boutons, however, do not have the time for proper development of pre– and postsynaptic structures, and are thus referred to as ghost boutons. Because expressing *Copia^pol^*-shRNA and *Copia^gag^*-siRNA cause larvae to have a larger number of synaptic boutons, we hypothesized that a saturating stimulus, such as the acute stimulation, may suppress the effect, and so we carried out subthreshold K^+^ stimulation (*20*). In wildtype larvae, five cycles of stimulation are required for maximal ghost bouton formation and three cycles are insufficient to induce a significant increase. Therefore, we used three cycles of K^+^ stimulation (subthreshold), which we verified did not induce ghost bouton formation in wildtype larvae (Fig. 3F-I).

However, subthreshold stimulation of larvae expressing *Copia^pol^*-shRNA or *Copia^gag^*-siRNA in presynaptic neurons induced significant ghost bouton formation similar to the levels found in wildtype larvae given five cycles of stimulation (Fig. 3J, S5G). Further, we found that presynaptic disruption of *Copia^gag^* induced a significant increase in quantal content (Fig. 3K), which indicates higher synaptic vesicle release and is consistent with increased acute plasticity.

#### Copia and dArc1 have an antagonistic relationship

We noticed that manipulating expression of *Copia* and *dArc1* at the *Drosophila* NMJ leads to opposite effects on neuronal plasticity. Both proteins appear to signal through the recently discovered ViSyToR pathway, which involves proteins forming viral-like capsids, binding to RNA transcripts (their own as well as others) and using EVs to transfer across the NMJ (*21*). We therefore investigated whether the two proteins have antagonistic interactions by examining the effects of genetic manipulation of one on the localization of the other. Expressing *Copia^pol^*-shRNA or *Copia^gag^*-siRNA presynaptically leads to a large accumulation of α-dArc1 signal both pre– and postsynaptically (Fig. 4A-C and 4F-G). We verified that the increased dArc1 protein level corresponded with an increase in its RNA transcript in both pre-and postsynaptic tissues (Fig. S5C-D). Likewise, expressing *dArc1*-shRNA in larval neurons resulted in an increase of α-Copia^gag^ signal in the pre– and postsynaptic compartments (Fig. 4D-E and 4H-I).

**Figure 4.**
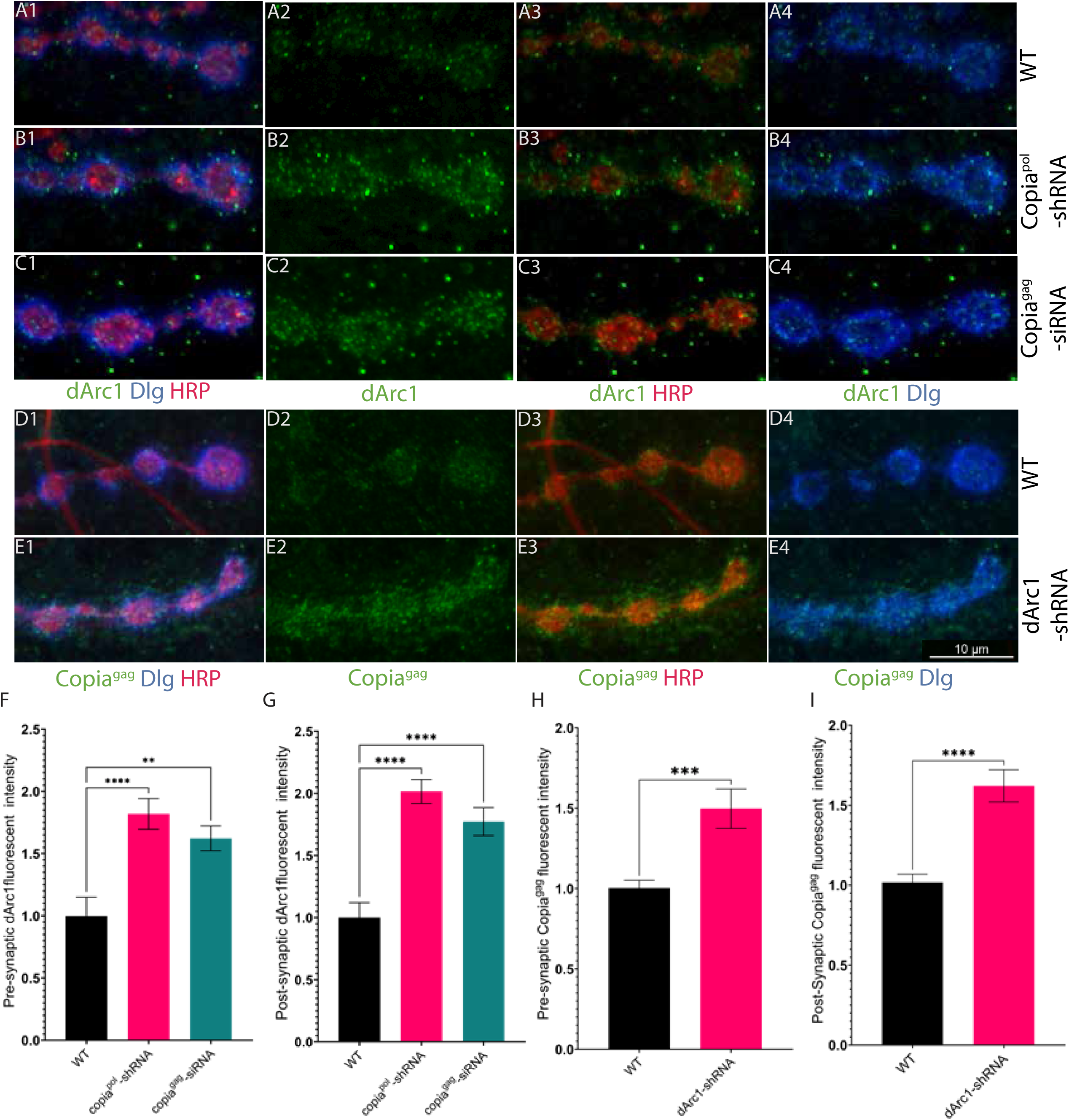
*Copia* and *dArc1* have an inverse relationship at the *Drosophila* NMJ. A–C. The presynaptic knockdown of *Copia* with *Copia*^pol^-shRNA (B1–B4) and *Copia*^gag^-siRNA (C1–C4) leads to a significant increase in α-dArc1 staining both pre– and postsynaptically at the larval NMJ compared to wildtype (A1–A4). D and E. The presynaptic knockdown of *dArc1* (E1– E4) leads to a significant increase in α-Copia^gag^ staining both pre– and postsynaptically at the larval NMJ compared to wildtype(D1–-D4) F–I. Quantification of the data represented in A–E. DLG = α-Discs Large (postsynaptic marker), HRP = α-horseradish peroxidase (presynaptic marker). N= number of animals/NMJs quantified in A) 8/14, B) 8/14, C) 8/14, D) 11/20 and E) 8/13. WT is C380-Gal4/Canton-S for panels A, D, and F-I. ns p≥0.05, * p<0.05, ** p<0.01, *** p<0.001, and **** p<0.0001.

#### *Copia* is genetically epistatic to *dArc1*

Because *dArc1* and *Copia* appear to have antagonistic effects on each other’s expression and induce opposite phenotypes, we tested whether they interact genetically by disrupting *Copia* in a *dArc1* null background. *dArc1* null mutants have substantially decreased bouton formation, which is consistent with our previous studies (Fig. 5A-C and 5F). Disrupting *Copia* expression in neurons in a *dArc1* null background results in a substantial increase in bouton numbers (Fig. 5A, 5D-F) as well as large increases in hyperbudding (Fig. 5D2-E2 and 5G). The mutually antagonistic relationship between expression and the result that *Copia* disruption reverses the *dArc1* phenotype shows an interaction between these two genes at larval NMJs and indicates that *Copia* is genetically epistatic to *dArc1* (Fig. 5H) (*22*).

**Figure 5.**
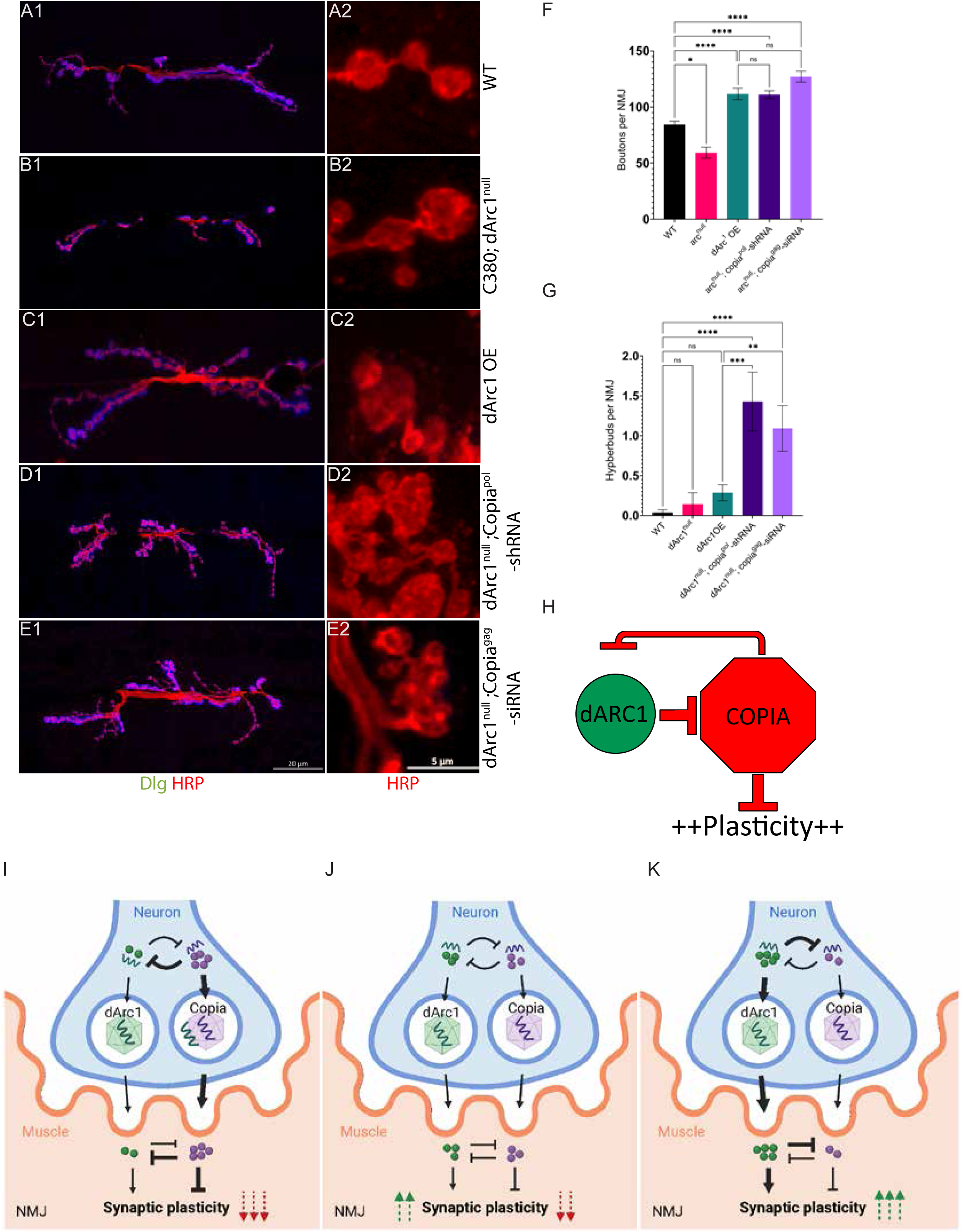
*Copia* is epistatic to *dArc1* in controlling synaptic plasticity. A–C. *dArc1* null (trans-heterozygous) flies (B1) show a substantial reduction in bouton formation compared to wildtype controls (A). C. There is a striking increase in bouton numbers (C1), but not hyperbudding (C2) in flies that are overexpressing *dArc1* presynaptically. D, E. Flies presynaptically expressing either *Copia*^pol^-shRNA (D) or *Copia*^gag^-siRNA (E) in an *dArc1* null background have increased number of boutons and hyberbudding. F, G. Comparison of bouton number and hyperbudding in wildtype, *dArc1* null, *dArc1* OE, and *dArc1* null expressing knockdown constructs against either *Copia*^pol^ or *Copia*^gag^ in neurons. H. The NMJ is in a state of high potential for plasticity and Copia represses plasticity in genetic epistasis to dArc1, as such removing both from the NMJ results in increased plasticity, consistent with our observations. I-K. Copia and dArc1 capsids compete for dArc1 mRNA. Illustrated in panel I, Copia capsids sequester *dArc1* mRNA this leading to a reduction in plasticity. While in K, a reduction of Copia binding to *dArc1* mRNA leads to an increase in dArc1 capsids and increase in plasticity, while in panel J, there is a balance of Copia and dArc1. DLG = α-Discs Large (postsynaptic marker), HRP = α-horseradish peroxidase (presynaptic marker). N= (by genotype from top to bottom; number of animals/NMJs quantified) 9/17, 7/12, 12/22, 4/8, 6/11 in (H) and (I). Full Genotypes in materials and methods. ns p≥0.05, * p<0.05, ** p<0.01, *** p<0.001, and **** p<0.0001.

## Discussion

In this work, we show evidence for the role of the TE *Copia* in the regulation of structural synaptic plasticity at the *Drosophila* larval NMJ. The splice variant *Copia^gag^* is enriched at the NMJ and likely transfers in a pre– to postsynaptic manner in EVs. Knockdown of *Copia^gag^* leads to an increase in the number of boutons, suggesting that *Copia* is an inhibitor of synapse formation. In larvae where *Copia* expression was disrupted, a subthreshold stimulation of potassium was sufficient to induce a significant change in activity-dependent bouton formation, which is consistent with the role of *Copia* in inhibiting activity-dependent plasticity. Additionally, we show that *Copia* is genetically epistatic to the master regulator of plasticity *dArc1* (Fig. 5H). Taken together, our observations uncover a novel physiological role for a fully functional TE at the NMJ, which acts as a potent regulator of structural synaptic plasticity.

How do Arc and *Copia* antagonize each other’s function at the NMJ? Being that Copia is epistatic to dArc1, at least in terms of structural synaptic plasticity at the larval NMJ, it suggests dArc1 acts as an inhibitor of Copia, itself an inhibitor of plasticity. However, this does not tell us how this interaction might be carried out at a molecular level. Because both form capsids using a related protein fold, one possibility is that each protein prevents assembly of the opposing capsid by binding to an assembly intermediate and blocking further oligomerization (*i.e.* a “Poison Pill” hypothesis). We disfavor this hypothesis due to the distinct interaction surfaces within the Copia and dArc capsids. A more likely possibility is that the capsids genetically antagonize the other at the level of cargo transport. We have observed that dArc1 protein does not precipitate Copia mRNA (Fig. S1F) (*5*), while Copia immunoprecipitates with both Copia mRNA and dArc1 mRNA (Fig. S1D and S1G). We also found that Copia^gag^ needs RNA to assemble into a capsid (Fig. S4). Taken together, the competition of dArc1 and Copia is likely at the state of mRNA binding. If Copia binds mRNA, it will more efficiently form capsids and sequester more dArc1 mRNA, presumably inhibiting dArc1 function (Fig. 5 I and 5J). In contrast, if dArc1 binds proportionally more dArc1 mRNA this would protect dArc1 from Copia sequestration, while also reducing RNA substrate for Copia to bind mRNA (Fig. 5K). Future experiments will investigate these possibilities. This latter model is testable, and as the association of dArc1 mRNA would be expected to change association with dArc1 or Copia, presumably in capsids, in different states of plasticity.

The *dArc1* gene likely arose from co-option of a TE by the host for a cellular function, referred to as domestication. It has been proposed that *Arc* was inserted under the control of a nearby neuronal transcriptional element, and that the expression of *Arc* in response to these elements influenced plasticity during development. This effect on plasticity was advantageous and selected for, which resulted in the gene remaining in the host genome. *Copia* is strikingly different because it is present as both a full-length and an alternatively spliced isoform. Disrupting expression of both the full-length and spliced transcripts causes a neuronal plasticity phenotype. It appears that the alternatively spliced isoform is more highly enriched in neuronal tissue than the full-length transcript, and knockdown of this specific isoform largely mimics knockdown of full-length *Copia*. This suggests an exciting new example of TE domestication that relies on the “host’s” alternative splicing system. Our data indicate that a spliced isoform of the viral-derived TE *Copia* is needed to properly regulate synaptic plasticity in antagonism to Arc, a TE-derived gene. We cannot determine if this is host selection for Copia-mediated synaptic plasticity, or if Copia is parasitizing the host. It is possible we are observing the very beginning of the domestication process or symbiosis of horizontally transferred viral-like genetic elements.

Recently, researchers engineered mammalian capsid-like protein, peg10, to take advantage of the ViSyToR pathway in order to transfer RNA cargo (*23*) and it is possible that other capsid-like genes can act in a viral-like manner. For instance, the retrotransposon *Cer1* in *C. elegans* has been shown to form capsids and may have a role in pathogen avoidance (*24*). These studies, along with our discovery that the TE *Copia* regulates synaptic plasticity, raise the possibility that eukaryotes have harnessed viral genomes to create capsid-like structures that regulate cellular functions. Further, these viral-like pathways could be repurposed for pharmaceutical uses such as improved delivery of gene therapies. The roles of TEs in synaptic development and other processes need to be further explored to better understand molecular underpinnings of cellular development and function as well as to understand the potential functions of a large part of eukaryotic genomes that have traditionally be thought of as “junk.”

## Figure legends

**Figure S1.**
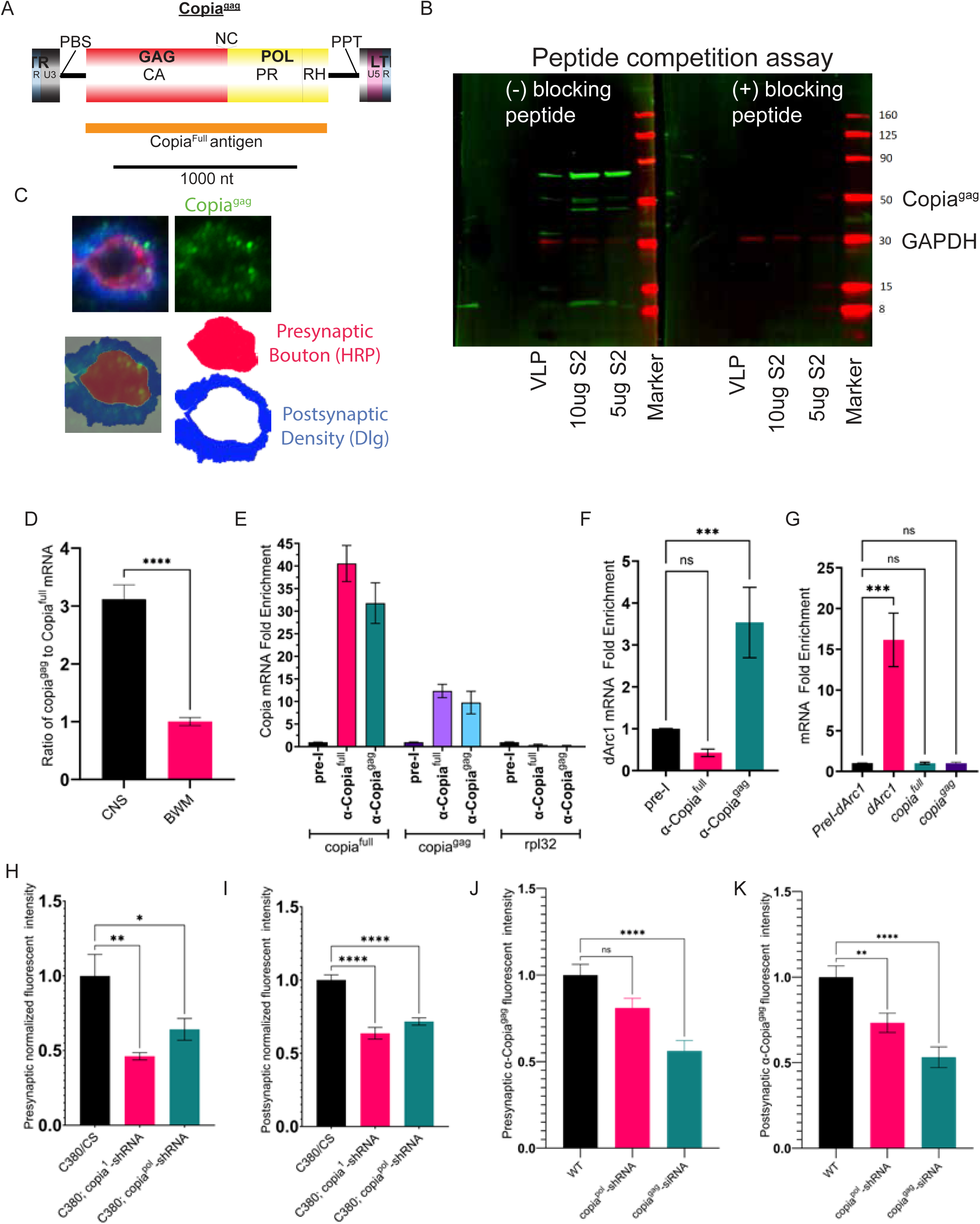
Copia protein and mRNA are enriched in Drosophila larval CNS. **A.** Schematic of Copia^gag^, the orange bar represents the region of Copia^gag^ used to generate a Copia^full^ antibody. B. Peptide competition assay, on the left lysates (labelled below) are probed with α-Copia^gag^, on the right, the blot is incubated with the Copia^gag^ antigen, Copia but not GAPDH staining is reduced. C. Diagrammatic representation of a bouton, highlighting the presynaptic bouton and postsynaptic density (HRP = Horseradish peroxidase antibody and DLG = discs large antibody). D. dPCR showing the ratio of Copia^gag^ to Copia^full^ transcript in the CNS vs. BWM, samples were normalized to Rpl32 transcript. E. *Copia* RNA immunoprecipitation (RIP) using Copia antibodies show that *Copia* mRNA is enriched in comparison to other mRNAs (BWM). F. Copia^gag^ immunoprecipitations enriches *dArc1* mRNA. G. dArc1 immunoprecipitation enriches *dArc1* mRNA but not *Copia^full^* nor *Copia^gag^* mRNA, all columns represent anti-dArc1 immunoprecipitation or pre-immune control, x-axis labels indicate RNA probes used respective column dPCR. H. *Copia^1^*-shRNA shows a similar reduction of presynaptic α-Copia^gag^ when expressed presynaptically (C380-Gal4), as compared to *Copia^pol^*-shRNA, as well there is a consistent decrease in postsynaptic α-Copia^gag^ (I) (For H and I, N= number of NMJs quantified WT (C380/CS) 8, C380; copia^1^-shRNA 7 and C380; copia^pol^-shRNA 7). J. Quantification of α-Copia^gag^ in the presynaptic compartment, with either Copia^pol^-shRNA or Copia^gag^-siRNA expressed presynaptically. K. Quantification of α-Copia^gag^ in the postsynaptic compartment, with either Copia^pol^-shRNA or Copia^gag^-siRNA expressed presynaptically, (N= from left to right; number of animals/boutons quantified, 8/24, 9/20, 11/20).

**Figure S2:**
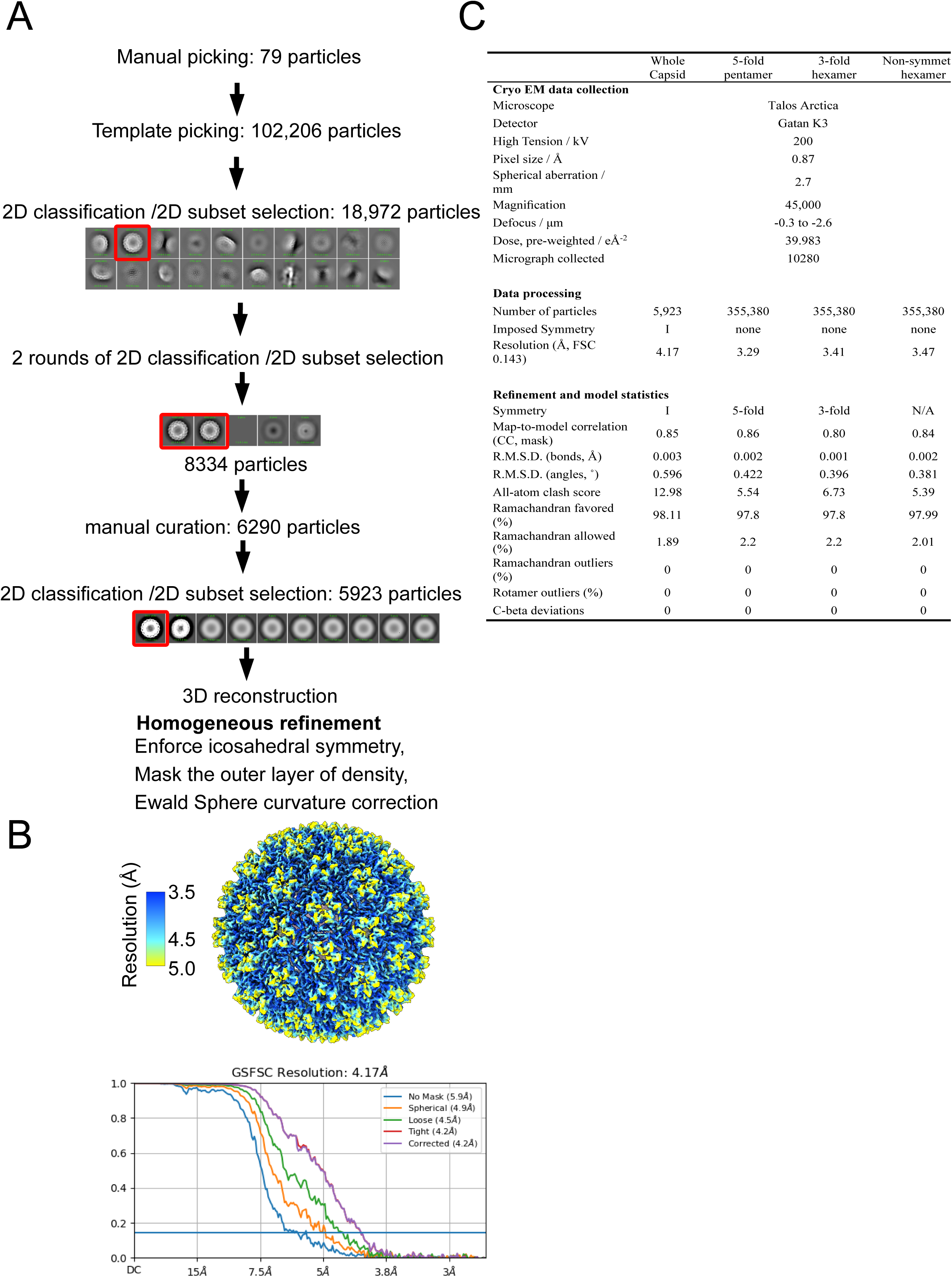
Cryo-EM structure determination of Copia^Gag^ capsid. All data processing was performed using cryoSPARC. A. Workflow for cryo-EM structure determination. Particles were first manually picked, which then were used for the template picker function. Particles were extracted and underwent three rounds of 2D classification, followed by manual curation and another round of 2D classification. We then performed 3D reconstruction with icosahedral symmetry enforced, and the outer layer of density masked. B. Local resolution and FSC curves of the cryo-EM map. Local resolution of the reconstructions and a representative section of each density map. The overall resolution of each map was determined by the FSC of each half-map using Gold-standard cutoff of 0.143. C. Table of Cryo-EM data collection, processing, and model statistics.

**Figure S3:**
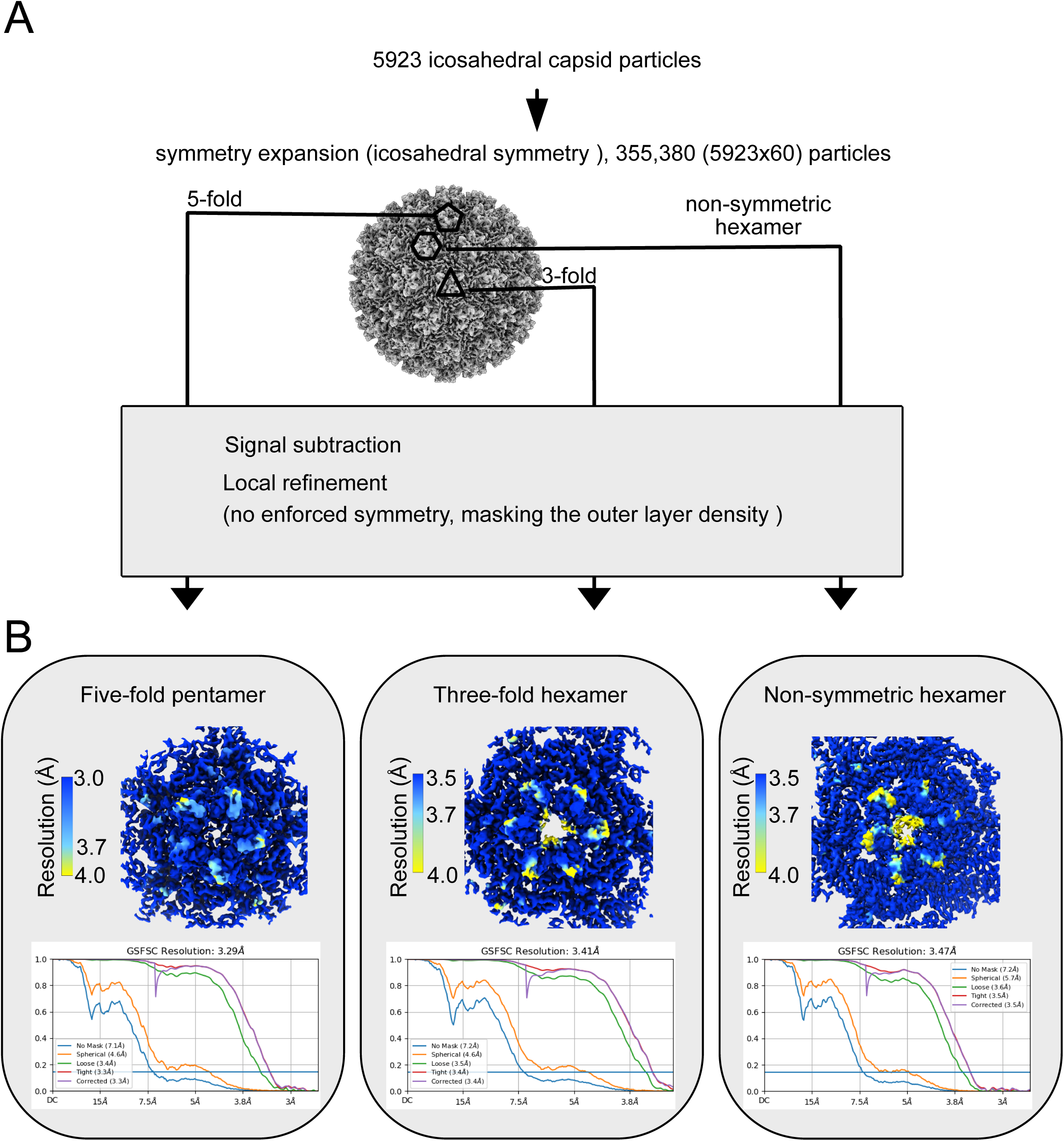
High resolution structure determination of Copia^Gag^ capsomers. All data processing was performed using cryoSPARC. A. Workflow for cryo-EM structure determination of individual capsomers. The capsid particle stack is identical to that of the full capsid (Fig. S2). Symmetry expansion was used to isolate individual capsomers. After signal subtraction and masking the outer layer, the reconstruction of each capsomer was refined locally. B. Local resolution and FSC curves of the cryo-EM maps. Local resolution (Fourier shell correlation (FSC)=0.143) of the reconstructions and a representative section of each density maps. The overall resolution of each map was determined by the FSC of each half-map using Gold-standard cutoff of 0.143.

**Figure S4:**
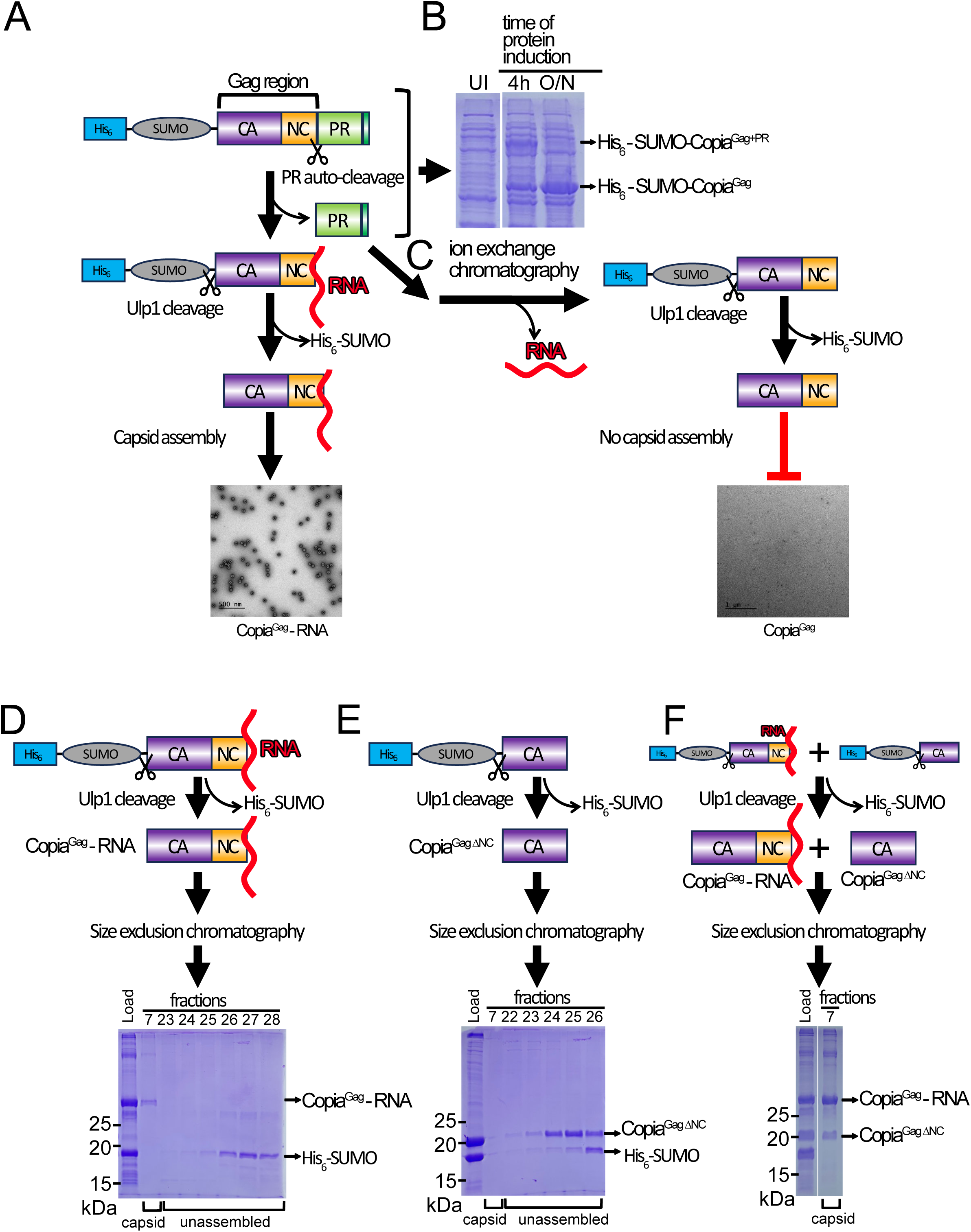
Assembly requirements for Copia^Gag^. A. Copia^Gag-PR^ auto-processes to cleave off the Protease (PR) region. Subsequent removal of the His_6_-SUMO tag triggers assembly into the capsid form. B. Auto-processing *in cellulo*. Uninduced *E. coli* cells (UI) show no expression of Copia^Gag+PR^. After 4 hours of expression at 18 °C, substantial full-length Copia^Gag+PR^ is observed, but after overnight expression nearly all Copia^Gag+PR^ is autoprocessed into Copia^Gag^ and Copia^PR^. C. Removal of RNA through ion exchange chromatography results in a Copia^Gag^ that does not assemble into capsids. D. Monitoring capsid assembly by Size Exclusion Chromatography. The capsid form of Copia^Gag^ elutes in fraction 7, while unassembled protein elutes in later fractions. E. A construct that lacks the RNA-binding Nucleocapsid domain (Copia^GagΔNC^) does not assemble into capsids. F. A mixture of Copia^Gag^ and Copia^GagΔNC^ results in both proteins assembling into capsids. This result illustrates that Copia^GagΔNC^ is assembly-competent, but lacks the ability to trigger assembly in isolation, presumably because of the lack of bound RNA.

**Figure S5.**
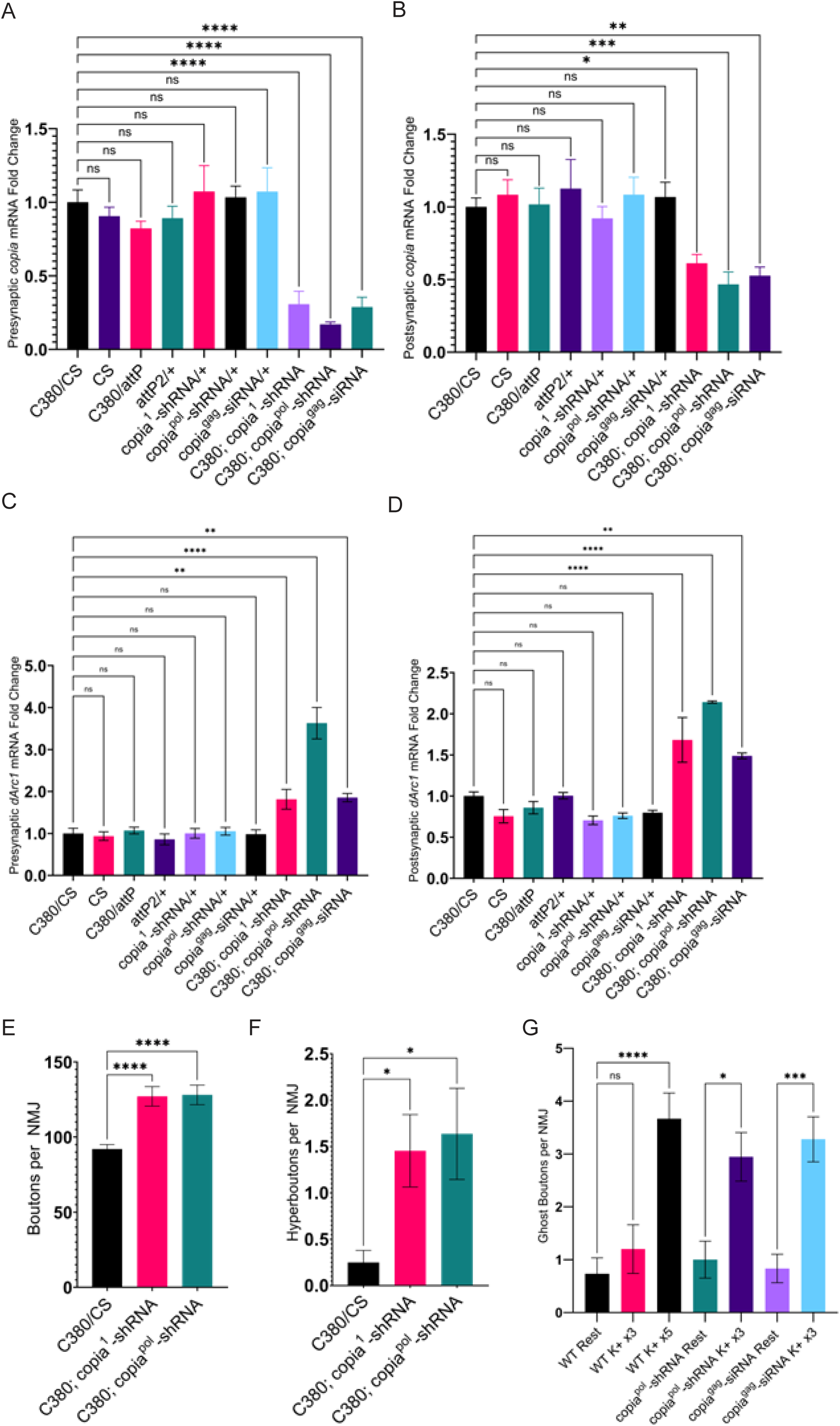
Copia and dArc1 mRNA have an inverse relationship at the Drosophila NMJ. A, B. Only in larvae expressing a *Copia* specific siRNA/shRNA presynaptically is there a significant decrease of Copia in either the pre– or postsynaptic compartments. C and D (N=3 for A-D). In flies expressing *Copia* si/shRNA *dArc1* mRNA expression is increased. E. *Copia^1^*-shRNA and *Copia^pol^*-shRNA driven presynaptically result in similar increases in bouton numbers and hyperbudding (F) (E and F, N=13 (C380/CS), 12 (C380; copia^1^-shRNA) and 11 (C380; copia^pol^-shRNA). G. Ghost boutons observed under various potassium (K+) conditions and mutant combinations.

## Methods

### Experimental Model and Subject Details

The following fly lines were used in generating the presented data: UAS-Copia^pol^-shRNA (see below), UAS-Copia^Gag^-siRNA (see below), UAS-dArc1-RNAi2 (*5*) w; dArc1^esm18^ (RRID:BDSC_37530, Bloomington *Drosophila* stock center, BDSC), y[1] w[67c23]; P[y[+t7.7]=CaryP]attP2 (RRID: RRID:BDSC_8622, BDSC), Canton-S (1, BDSC), C380-Gal4 (*25*) and C57-Gal4 (*25*). Female third-instar larvae were used for all NMJ dissections.

### Fly Husbandry

All flies were raised on low yeast molasses formulation *Drosophila* food at either 25°C or 29°C (Gal4/RNAi crosses).

### Addressing Copia-Mediated Off-target Effects

It has been proposed that TEs regulate physiological functions through intermediate chimeras, whereby TEs co-splice with nearby genes (*26*). If a fragment of the *Copia* transcript is co-spliced with another transcript, then repressing *Copia* may affect NMJ development and/or function through functioning in a chimeric intermediate. We have taken several steps to investigate this possibility. As with other RNAi constructs directed against a canonical gene, we created several RNAi constructs that were designed to target *Copia* specifically. While making multiple RNAi constructs targeting different regions of *Copia* (Fig 1B), we searched for fragments of *Copia* in the annotated *Drosophila* genome and found 38 full-length copies, which is consistent with previous analysis (*27*). We searched the entire genome for regions that contain either Copia^1^-shRNA or Copia^pol^-shRNA and found matches to both RNAi constructs in each of the 38 copies as well as another 21 matches to fragments of *Copia* scattered throughout the genome. We were able to only search within these fragments for matches to Copia^gag^-siRNA because this RNAi construct is too short to effectively search for in the entire genome. No mature mRNA transcripts contained appreciable sequence matches either Copia^1^-shRNA or Copia^pol^-shRNA. While we did successfully identify chimeric transcripts, we did not find any that were targeted by the RNAi constructs that we used to disrupt *Copia* expression and being that these sequences are not near known seeds of *Copia*, this suggests that these chimeras are a product of deep sequencing noise (*28*).

### Constructs

For *Copia^pol^*-shRNA and other dsRNA constructs, the insert was synthesized (see attached construct sequence file), then cloned into pwallium10-roe as described in Ni et al 2011 (*29*). For the *Copia^gag^* siRNA, the forward and reverse primers were synthesized (see attached construct sequence file), annealed, and then cloned into pwallium20 as described in Ni et al 2011(*29*).

### Immunocytochemistry and Antibodies

*Drosophila melanogaster* third instar larva body wall muscles were dissected in calcium-free saline and fixed in either Bouin’s fixative (0.9% (v/v) picric acid, 5% (v/v) glacial acetic acid, 9% (w/v) formaldehyde) or 4% (w/v) paraformaldehyde (PFA) in 0.1M phosphate buffer, pH 7.2. Fixed samples were washed and permeabilized in phosphate-buffered saline (PBS) with Triton X-100 (PBT; 0.1M phosphate buffer; 0.2% (v/v) Triton X-100) and incubated in a primary antibody overnight at 4°C. The samples were then washed three times with PBT, incubated with secondary antibodies for two hours at room temperature, washed three times, and then mounted in Vectashield Hardset Mounting Media (Vector Laboratories Inc.). The following antibodies were used: rabbit anti-Copia^Full^, 1:1000 (see below), rabbit anti-Copia^Gag^, 1:5000 (see below), rabbit anti-dArc1, 1:500 (*30*), rabbit anti-Discs-large (DLG), 1:40,000 (*31*), and mouse anti-DLG, 1:200 (Developmental Studies Hybridoma Bank (DSHB), 4F3). DyLight-conjugated and Alexa Fluor-conjugated secondary antibodies were obtained from Jackson ImmunoResearch (DyLight-405-conjugated goat anti-horseradish peroxidase (HRP), Alexa Fluor-594-congugated goat anti-HRP, Alexa Fluor-488-congugated donkey anti-Rabbit, Alexa Fluor-594-congugated goat anti-rabbit, Alexa Fluor-647-congugated goat anti-mouse) and were used at 1:200, as described above.

Copia^Full^ antibodies were generated against a Copia antigen (***see*** ***Figure 1***) by immunizing rabbits with the entire Copia^gag^ protein (Pocono Rabbit Farm and Laboratory), while the Copia^gag^ antibodies were generated against a Copia peptide antigen (***see*** ***Figure 1***) by immunizing rabbits with the peptide LMVVKNSENQLADIC (GenScript).

### Activity Paradigm

Potassium stimulations were carried out as described in Ataman et al 2008 (*20*). Larva were dissected in low-calcium (0.1mM) HL3 saline(*32*), then pulsed with a series of high potassium (90mM) saline. Each pulse was spaced out by a 15-minute rest period in low-calcium HL3 saline. The 5-cycle potassium stimulation consisted of three 2-minute pulses, one 4-minute pulse, and one 6-minute pulse, which was then followed by a final 15-minute rest. The 3-cycle (subthreshold) potassium stimulation consisted of three 2-minute pulses followed by a 15-minute rest period. Following the 90-minute pulse-rest cycle, samples were fixed with 4% PFA and processed for immunocytochemistry as described above.

### Confocal Microscopy and Signal Intensity Measurements

Z-stacked images were acquired using a Zeiss LSM 800 confocal microscope equipped with a Zeiss 63X Plan-Apochromat 1.40 NA DIC M27 oil immersion objective and a Zeiss 40× Plan-Apochromat 1.30 NA DIC (UV) VIS-IR M27 oil immersion objective (Zeiss Inc). After image acquisition with identical settings, the images were quantified as previously described (*33*). In brief, volumetric measurements of the boutons of interest bound by HRP staining, a nervous system-specific marker that delineates the presynaptic volume, was selected in parallel to the postsynaptic volume bound by the DLG (a subsynaptic reticula mark) signal (Fig. S1C, cartoon). Fluorescence intensity of target antibodies was measured, and the intensity was normalized to the bouton volume calculated from the HRP and DLG signal and all data were normalized to wildtype values. Quantitations were made using the Volocity Version 6.3 software (Quorum Technologies Inc).

We found that the knockdown of *dArc1* caused the α-Copia^gag^ signal to become saturated when capturing images at laser intensity and detector sensitivity that were needed for dynamic intensity in the wildtype flies. To collect data in a linear range of the signal intensity, we normalized the data to the *dArc1*-siRNA values rather than wildtype so that a shorter exposure could be taken. Total fluorescent intensity was measured in the larvae given subthreshold stimulation of potassium.

### RNA Immunoprecipitation

Wildtype *Drosophila* third-instar larvae were dissected, and the CNS and BWM were collected in separate tubes containing RIPA buffer (Abcam) supplemented with protease inhibitors (Roche) and RNase inhibitor (Invitrogen) as previously described (*30*). Similarly, S2 cells were grown to confluency, washed with ice-cold DPBS (Sigma) and resuspended in RIPA buffer. Tissue and cell lysates were homogenized using 0.5mm glass beads at 4°C using a Bullet Blender 24 Gold homogenizer (Next Advance Inc.). Lysates were then centrifuged at 4°C to remove cell debris. Supernatants were precleared against Protein A/G magnetic beads (Pierce), and then incubated overnight at 4°C with either anti-Copia^Full^, anti-copia^gag^, anti-dArc1 antibodies, or equal amounts of pre-immune serum. Samples were then incubated for 2 hours at 4°C with protein A/G magnetic beads and washed several times with RIPA buffer. For immunoblotting, beads were incubated directly with 4X protein loading buffer (Li-Cor) with 2-Mercaptoethanol (Sigma). For digital PCR, RNA was eluted from the beads with RLT buffer (QIAGEN) supplemented with 2-mercaptoethanol and then purified using the RNeasy mini kit (QIAGEN) for the QIAcube connect (QIAGEN) with DNase digest using RNase-free DNase set (QIAGEN).

### Digital PCR (dPCR)

RNA samples were reverse transcribed into cDNA using the Superscript IV first-strand synthesis reaction (Invitrogen) following the manufacturer’s protocol with RNase H digest. The dPCRs were multiplexed in 26K 24-well or 8.5K 96-well QIAcuity nanoplates (QIAGEN) using a QIAcuity system (QIAGEN). For the reactions, either QIAcuity EvaGreen master mix or probe master mix (QIAGEN) were used with the gene-specific primer sets for *dArc1*, *Copia*^Full^, *Copia*^gag^, Rpl32, and 18S rRNA or their probes (ThermoFisher and/or IDT). Data were processed in the QIAcuity Software Suite (QIAGEN) where absolute values (copies/µL) were obtained, and normalized expression was derived.

### Immunoprecipitation

Third-instar wild type *Drosophila* larvae were dissected in ice-cold Ca^2+^-free saline, and CNS and BWM were collected and homogenized as described above. S2 cells were grown to confluency, washed in ice-cold DPBS (Sigma), resuspended in RIPA buffer (Abcam) supplemented in protease inhibitor cocktails (Roche), and homogenized as described above. Lysates were centrifuged at maximum speed at 4°C for 10 minutes. Protein concentration was determined by Qubit protein assay (Invitrogen) in a Qubit 4 fluorometer (Invitrogen).

Supernatants were incubated with the respective antibody overnight at 4°C with gentle rotation, then the protein-antibody complexes were incubated with protein A/G magnetic beads (Pierce) for an hour at room temperature. The samples were then washed with buffer several times, and then a final time in distilled water. The magnetic beads were eluted with protein sample buffer at room temperature for 10 minutes with gentle rotation or boiled at 95°C for 10 minutes.

### Western Blotting

Immune complexes from RIP and IP experiments were incubated at room temperature or 95°C for 10 min. proteins were separated in Mini-Protean TGX stain-free 4–20% precast gels (Bio-Rad) under reducing and denaturing conditions. Proteins were transferred to an Immuno-Blot LF PVDF membrane (Bio-Rad) on a semi-dry Trans-Blot Turbo transfer system (Bio-Rad), blocked in Intercept blocking buffer (Li-Cor) and incubated with primary antibodies diluted in Intercept antibody diluent (Li-Cor) overnight at 4°C. Blots were washed, incubated with IRDye secondary antibodies (Li-Cor), washed again and finally imaged on a Li-Cor odyssey CLx imaging system.

### Electrophysiology

All experiments were performed on wandering third instar larvae raised at 29°C. Larvae were dissected under ice-cold hemolymph-like (HL-3.1) saline (*34*)containing 0.3 mM calcium. Body-wall muscles were visualized under a Zeiss Examiner D.1 using a 10× long-working distance objective paired with a 4X magnifier and continually superfused with HL-3.1 saline containing 0.5 mM calcium at RT. Recordings were done by impaling body-wall muscle 6-7 in abdominal segment 3 with a 20-25 MÙ electrode and were amplified using an Axoclamp 900A amplifier (Molecular Devices, San Jose, CA). Recordings were exported to a computer using an Axon Digidata 1550B (Molecular Devices, San Jose, CA). Data were analyzed using both Easy Electrophysiology and Origin software (OriginLab, Northampton, MA). Evoked excitatory junctional potentials (EJPs) were induced through application of a suction electrode (A-M Systems, Sequim, WA) to a cut segmental nerve and stimulated with a 1 ms suprathreshold stimulus at 0.5 Hz by an A365 stimulus isolator (World Precision Instruments, Sarasota, FL). Only muscle cells with a resting potential more negative than –60mV mV were used for analysis, and at least five cells were analyzed for each genotype. Data was compiled in GraphPad Prism version 9.5.0 (GraphPad Software)., outliers removed, and statistical analysis as outlined below.

### Expression, purification, and assembly of Copia^Gag^ capsids

The short form of Copia encodes Copia^Gag^ and the protease domain (PR), which is termed Copia^Gag+PR^ in this study. In order to test if Copia^Gag+PR^ can form capsid like particle through Copia^Gag^ self-assembly, we expressed and purified the Copia^Gag+PR^ protein using the *E. coli* expression system. Copia^Gag+PR^ was fused to an N-terminal His_6_-SUMO tag and cloned into the pSMT3 vector. pSMT3_His_6_-SUMO-Copia^Gag+PR^ was transformed into *E. coli* BLR (DE3) cells for protein expression. *E. coli* cells harboring pMST3_His_6_-SUMO-Copia^Gag+PR^ were cultured in 1 L Terrific Broth (containing 30 mg/L Kanamycin) at 37°C with shaking. When cells reached an optical density (600 nm) of 0.6 – 0.8, the culture was induced by adding isopropyl β-D-1-thiogalactopyranoside (IPTG) to the final concentration of 0.2 mM. The induction was done at 18 °C with shaking for 24 hours. After induction, cells were harvested by centrifuging at 5000xg for 20min at 4 °C.

During the induction, the protease domain in Copia^Gag+PR^ auto-processes the His_6_-SUMO-Copia^Gag+PR^ polyprotein to generate His_6_-SUMO-Copia^Gag^ (Fig. S4). His_6_-SUMO-Copia^Gag^ was purified using Ni-affinity chromatography. All the following purification steps were performed at 4°C. The cell pellet was resuspended in buffer A (25 mM HEPES, pH7.4, 150 mM NaCl, 10% Glycerol, 100 µM ZnCl_2_, 2 mM 2-mercaptoethanol) supplemented with 20 mM imidazole and protease inhibitors. Cells were lysed by high pressure cell disruption on ice. Cell lysate was clarified by centrifuging at 20,000 ×g for 40 min at 4°C. The supernatant was transferred and filtered through a 0.45 µm pore size PVDF membrane before loaded onto the Ni-affinity column (Cytiva, HisTrap HP Cat#17524801). His_6_-SUMO-Copia^Gag^ was eluted off the Ni-affinity column through a gradient of 20-500 mM imidazole. Only elution fractions with imidazole concentration from 400-500 mM were pooled and digested with His_6_-tagged SUMO-protease Ulp1 to cleave off the N-terminal His_6_-SUMO tag and trigger Copia^Gag^ capsid assembly. Copia^Gag^ capsids were further purified by removing free His_6_-SUMO and Ulp1-His_6_ by passing the protein across the Ni-affinity column.

The His_6_-SUMO-Copia^GagΔNC^ construct was generated by mutating residue H186 to a stop codon in the wild-type pSMT3_His_6_-SUMO-Copia^Gag+PR^ plasmid using Quikchange mutagenesis. His_6_-SUMO-Copia^GagΔNC^ was induced and Copia^GagΔNC^ was purified the same way as the wild-type Copia^Gag^ described above.

### Copia^Gag^ protein RNA depletion

To remove the RNA from the Copia^Gag^ protein purified in Fig. S4, Ni-affinity purification was first performed as described above. After Ni affinity purification, the pooled protein fractions were immediately loaded onto an anion exchange column (Mono Q™ 5/50 GL, Cytiva # 17516601), and protein was eluted off the column through a gradient of 150-1000 mM NaCl supplemented in buffer 25 mM HEPES, pH7.4, 10% Glycerol, 100 µM ZnCl_2_, 2 mM 2-mercaptoethanol. Fractions with pure His_6_-SUMO-Copia^Gag^ were pooled and dialyzed against buffer 25 mM HEPES, pH7.4, 600mM NaCl, 10% Glycerol, 100 µM ZnCl_2_, 2 mM 2-mercaptoethanol at 4°C overnight. The His_6_-SUMO tag was then cleaved off by Ulp1 protease and removed by Ni-affinity chromatography. The purified protein Copia^Gag^ (RNA free, A260/280 = 0.6) was dialyzed against storage buffer 25 mM HEPES, pH7.4, 600mM NaCl, 20% Glycerol, 100 µM ZnCl_2_, 2 mM 2-mercaptoethanol, flash frozen in liquid nitrogen and stored at –80°C.

To monitor capsid assembly by size exclusion (as shown in Fig. S4 D-F), Ulp1 protease was added to the purified His_6_-SUMO-Copia^Gag^/ His_6_-SUMO-Copia^GagΔNC^ protein after Ni-affinity purification to cleave off the His_6_-SUMO. After overnight Ulp1 digestion, the sample was loaded onto the gel filtration column (Superose™ 6 Increase 10/300 GL, Cytiva # 29091596) and eluted in buffer 25 mM HEPES, pH7.4, 150mM NaCl, 10% Glycerol, 100 µM ZnCl_2_, 500 mM Imidazole, 2 mM 2-mercaptoethanol. Assembled capsids elute at fraction 7 (retention volume = 7 mL), while the unassembled Copia^GagΔNC^ protein at fraction 24 to 26 (retention volume = from 17 to 18 mL). In Figure S4F, Ulp1 protease was added after mixing His_6_-SUMO-Copia^Gag^ and His_6_-SUMO-Copia^GagΔNC^. After overnight Ulp1 digestion, the protein mixture was loaded onto the gel filtration column (as described above), and capsids were eluted off the column at fraction 7.

### Negative-stain Transmission Electron Microscopy

For samples in Fig. S4A, Copia^Gag^ capsid protein was concentrated using a 100kDa MWCO Amicon® Ultra-15 Centrifugal Filter Unit(EMD Millipore) to a final concentration of 0.8 mg/ml. Protein concentration was determined by Bradford assay. Copia^Gag^ capsid protein was filtered by 0.22 µm pore size PVDF membrane filter prior to deposition on the grid for EM imaging.

Copper grids coated with carbon film (Electron Microscopy Sciences, CF200-Cu-50) were glow discharged on a PELCO easiGlow (Ted Pella) at 25 mA for 35 s (negative polarity) before use. 7 μl of sample was applied to the grid and incubated for 1 min. Excess sample was blotted on filter paper, then the grid was rinsed with 15 µl water (filtered by 0.22 µm pore size PVDF membrane) three times followed by staining with 1% uranyl acetate (pH 4.5) for 1 min and then blotted dry. Samples were imaged with a FEI Tecnai Spirit 12 transmission electron microscope at 120 kV equipped with a Gatan 4K camera.

For samples in Fig. S4C, Copia^Gag^ (RNA free, A260/A280 = 0.6) capsid protein was concentrated using a 100kDa MWCO Amicon® Ultra-15 Centrifugal Filter Unit (EMD Millipore) to a final concentration of 1.4 mg/ml. Protein was diluted 4-fold with buffer 25 mM HEPES, pH7.4, 10% Glycerol, 100 µM ZnCl_2_, 2 mM 2-mercaptoethanol and filtered by 0.22 µm pore size PVDF membrane. The capsid formation was assessed and imaged using negative-staining EM as described above for samples in Fig. S4A.

### Cryo-EM specimen preparation

Copia^Gag^ capsid protein was dialyzed against Cryo-EM buffer (25 mM HEPES, pH7.4, 150 mM NaCl, 100 µM ZnCl_2_, 2 mM 2-mercaptoethanol) overnight at 4°C. The protein sample was concentrated using 100kDa MWCO Amicon® Ultra-15 Centrifugal Filter Unit (EMD Millipore) to a final concentration of ∼0.8mg/ml and filtered by 0.22 µm pore size PVDF membrane prior to deposition on the grid for EM imaging.

Grids were washed by ethyl acetate and glow discharged on a PELCO easiGlow (Ted Pella) at 25 mA for 35 s (negative polarity) before use. 3.5 μL of Copia^Gag^ capsid protein sample was applied to a 300-mesh copper grid with continuous 2 nm carbon film (Electron Microscopy Sciences, QUANTIFOIL® C2-C15nCu30-50) at 10 °C with 90% humidity in a Vitrobot Mark IV (FEI). Sample was blotted from both sides for 8 s with blot force 0, then immediately vitrified by plunging into liquid ethane.

### Data collection

Micrographs were collected on a 200 kV Talos-Arctica electron microscope (FEI) equipped with a K3 Summit direct electron detector (Gatan). Images were collected at a magnification of 45,000x in counting mode with an unbinned pixel size of 0.87 Å and a total dose of 39.98 e/Å^2^ per micrograph, with a target defocus range of −0.3 to −2.9 μm. In total, 10280 micrographs were collected.

### Data processing

Copia^Gag^ capsid image processing and reconstruction are summarized in Supplementary Fig. S2 and S3. Dataset parameters are presented in Fig. S2B. All the reconstruction steps were done within the CryoSPARC package (*35*). Acquired micrograph frames were imported and motion corrected using patch-based motion correction followed by Contrast Transfer Function (CTF) estimation using patch CTF in CryoSPARC. Particles were picked using the template picker function. The template was generated by a training dataset of manually picked ∼80 particles.

A total of 102,206 particles were automatically picked using template picking and extracted into 896×896 pixel boxes (0.87 Å / pixel). Particles were binned to 224×224 pixel boxes (3.48 Å / pixel) during the extraction. Three rounds of 2D classification were performed to remove bad particles, resulting in a stack of 8334 particles. We manually curated the 8334, resulting in a stack of 6290 particles. After one more round of 2D classification, 5923 (94%) particles displayed icosahedral capsid structure and were selected as the final particle stack for further 3D reconstruction. The initial model was generated by *ab-initio* reconstruction with no symmetry enforced. The *ab-initio* model was further used in homogeneous refinement of the whole capsid with enforced Icosahedral symmetry (I) and Ewald Sphere curvature correction. The effective resolutions of the cryo-EM density maps were estimated by Fourier shell correlation (FSC = 0.143) between the two half maps (Fig. S2 and S3).

To further improve the resolution, we performed symmetry expansion as implemented in CryoSPARC. Local refinement at the five-fold pentamer, the three-fold hexamer and the non-symmetry related hexamer within the asymmetric unit was performed using symmetry expanded particles with the rest of the density subtracted. Local refinement was performed with no symmetry applied, and we determined the structures with resolutions of 3.29 Å for the five-fold pentamer, 3.41 Å for the three-fold hexamer and 3.47 Å for the non-symmetric hexamer (Fig. S3).

### Model building

The structures of Copia^Gag^ capsid (CA domain: residues 1 through 186) were determined by model building into the locally refined maps of the five-fold pentamer, three-fold hexamer, and the non-symmetric hexamer. No structures were determined for the inner layer that is likely corresponding to the NC domain (residue 186-270) and the associated nucleic acid, due to its disordered nature.

The initial model of Copia^Gag_NTD^ (residue1-90) and Copia^Gag_CTD^ (residue 91-186) were predicted using AlphaFold2 (*36*) which were separately fit as rigid bodies into the map density. The initial model was built using Coot (*37*) and refined in Phenix (*38*) using real-space refinement with rotamer, Ramachandran, and secondary structure restraints. For the five-fold and three-fold capsomers, symmetry constraints were applied during the refinement. The refinement and validation statistics were gained from the refinement report on Phenix and are listed in Fig. S2B. ChimeraX was used for molecular visualization and analysis (*39*).

### Isolation and Quantification of EVs from S2 Cells

Extracellular vesicles (EVs) were isolated from S2 cells cultured in serum-free medium at 22°C. The media was collected and first centrifuged at 500 × g for 5 min to pellet the cells. The supernatant was then collected and centrifuged at 2,000 × g for 10 min at 4°C to eliminate cell debris. To retain small EVs, the samples were further centrifuged at 10,000 × g for 30 min at 4°C. The supernatants were filtered with a 0.22 µm PES filtration unit (EMD Millipore), the samples were concentrated with a Centricon centrifugal filter (EMD Millipore) and the EV-rich filtride recovered. The samples were then purified by size exclusion chromatography using qEV isolation columns (Izon Sciences) on an automatic fraction collector (Izon Science). The fractions were then pooled based on protein concentration and Amicon centrifugal filters (EMD Millipore) were used to concentrate the samples. Samples were diluted in PBS and analyzed for size and concentration using the Exoid TRPS system (Izon Science). Samples that were not immediately used were stored at –80°C.

### S2 Cells Derived EVs Immuno-Electron Microscopy

Samples were prepared as previously described (*30*). In brief, the EV preparations were fixed overnight at 4°C in 2% PFA (EMS) and 5μL of sample applied to formvar coated gold grids (EMS) and incubated for 5 min. Grids were wicked on Whatman #50 filter paper (GE Healthcare) after which they were washed in 100mM Tris followed by additional washes in 100mM Tris + 50mM Glycine. Grids were blocked for 10 min in blocking buffer (EMS) and either incubated in Tris (control) or lysed with 0.05% saponin in Tris. After washing in Tris, they were incubated in primary antibody for 1hr, washed in Tris, and then incubated with anti-rabbit and/or anti-mouse conjugated to 10nm, 15nm or 18nm gold secondary antibodies (EMS). Grids were washed, post fixed with 1% glutaraldehyde, washed in water, and finally negative stained with 1% uranyl acetate for 30 s. Grids were imaged on an FEI Tecnai 12 Spirit equipped with a Gatan 4K camera.

### Quantification and Statistical Analysis

Statistical analyses for single comparisons were performed using a Student’s t test while comparison among multiple groups was done using a one-way analysis of variance (ANOVA) with the appropriate post hoc test. Statistical significance is indicated as ∗, p < 0.05; ∗∗, p < 0.001; ∗∗∗, p < 0.0001. Raw data files were processed with Excel (Microsoft) and data analysis for statistical significance was done with GraphPad Prism version 9.5.0 (GraphPad Software).

## Supporting information

Probes and Primers

