## Supplementary material for "Dueling Endogenous Viral-Like Sequences Control Synaptic Plasticity": Probes and Primers

| | | | |
|---|---|---|---|
| Copia forward (qPCR) | GGGATCAGGCAACCCGAATGAG | | IDT |
| Copia reverse (qPCR) | CTATTATCCTCTTCATTATAGGATAT | | IDT |
| Copia Full FWD | AGACTGCAATACCCCATGCC | | IDT |
| Copia Full REV | ATCTGGGCGTGTACAAAGCA | | IDT |
| Copia Gag FWD | GGTTGTCAAAAATTCAGAGAATCAAC | | IDT |
| Copia Gag REV | TGTCTCGTAACTCCACAAATCTC | | IDT |
| Copia Full Probe | AGCGAGATGATCACCTGAATG | GCCAGAGAGCAAGTTCAGAATA | IDT and Thermofisher |
| | GTGCTCTGCTGTTTCACTTTC | TCCTACAGAGAATCAACTGGCTGACA | |
| | AAGGGATCAGGCAACCCGAATGAG | TGTCTCGTAACTCCACAAATCTC | |
| Copia Gag Probe | AAAAGCGGTGTAACCATTTCGAAAA | GGTTGTCAAAAATTCAGAGAATCAAC | IDT and Thermofisher |
| | AGCAGGCAACGGTTTTGTAAATATG | TGCTGCGAGATTTGTGGAGTTACGA | |
| | CAGTTGATTCTCTGAATTTT | AGCATTCGATTGGTCGTCTT | |
| dArc1 Taqman Assay | Dm01823981_s1 | | ThermoFisher |
| Rpl32 probe | TGGTTTCCGGCAAGCTTCAA | | IDT |
| | TGTTGTCGATACCCTTGGGC | | IDT |
| | TCCGCCCAGCATACAGGCCCA | | IDT |
| Rpl32 Taqman Assay | Dm02151827_g1 | | ThermoFisher |
| Actin5C Taqman Assay | Dm02361909_s1 | | ThermoFisher |
| GAPDH Taqman Assay | Dm01843827_s1 | | ThermoFisher |
| | | | IDT |
